# A collaborative approach to improve representation in viral genomic surveillance

**DOI:** 10.1101/2022.10.19.512816

**Authors:** Paul Y. Kim, Audrey Y. Kim, Jamie J. Newman, Eleonora Cella, Thomas C. Bishop, Peter J. Huwe, Olga N. Uchakina, Robert J. McKallip, Vance L. Mack, Marnie P. Hill, Ifedayo Victor Ogungbe, Olawale Adeyinka, Samuel Jones, Gregory Ware, Jennifer Carroll, Jarrod F. Sawyer, Kenneth H. Densmore, Michael Foster, Lescia Valmond, John Thomas, Taj Azarian, Krista Queen, Jeremy P. Kamil

## Abstract

The lack of routine viral genomic surveillance delayed the initial detection of SARS-CoV-2, allowing the virus to spread unfettered at the outset of the U.S. epidemic. Over subsequent months, poor surveillance enabled variants to emerge unnoticed. Against this backdrop, long-standing social and racial inequities have contributed to a greater burden of cases and deaths among minority groups. To begin to address these problems, we developed a new variant surveillance model geared toward building microbial genome sequencing capacity at universities in or near rural areas and engaging the participation of their local communities. The resulting genomic surveillance network has generated more than 1,000 SARS-CoV-2 genomes to date, including the first confirmed case in northeast Louisiana of Omicron, and the first and sixth confirmed cases in Georgia of the emergent BA.2.75 and BQ.1.1 variants, respectively. In agreement with other studies, significantly higher viral gene copy numbers were observed in Delta variant samples compared to those from Omicron BA.1 variant infections, and lower copy numbers were seen in asymptomatic infections relative to symptomatic ones. Collectively, the results and outcomes from our collaborative work demonstrate that establishing genomic surveillance capacity at smaller academic institutions in rural areas and fostering relationships between academic teams and local health clinics represent a robust pathway to improve pandemic readiness.

**Author summary:** Genomic surveillance involves decoding a pathogen’s genetic code to track its spread and evolution. During the pandemic, genomic surveillance programs around the world provided valuable data to scientists, doctors, and public health officials. Knowing the complete SARS-CoV-2 genome has helped detect the emergence of new variants, including ones that are more transmissible or cause more severe disease, and has supported the development of diagnostics, vaccines, and therapeutics. The impact of genomic surveillance on public health depends on representative sampling that accurately reflects the diversity and distribution of populations, as well as rapid turnaround time from sampling to data sharing. After a slow start, SARS-CoV-2 genomic surveillance in the United States grew exponentially. Despite this, many rural regions and ethnic minorities remain poorly represented, leaving significant gaps in the data that informs public health responses. To address this problem, we formed a network of universities and clinics in Louisiana, Georgia, and Mississippi with the goal of increasing SARS-CoV-2 sequencing volume, representation, and equity. Our results demonstrate the advantages of rapidly sequencing pathogens in the same communities where the cases occur and present a model that leverages existing academic and clinical infrastructure for a powerful decentralized genomic surveillance system.

## Introduction

In late 2019, a newly emergent betacoronavirus, severe acute respiratory syndrome coronavirus 2 (SARS-CoV-2), was identified as the etiological agent associated with an outbreak of viral pneumonia in Wuhan, China [1]. The outbreak quickly grew to become a global public health emergency, now commonly referred to as the COVID-19 pandemic. Between November 2020 and January 2021, the first variants of concern (VOC) Alpha, Beta, and Gamma, were detected in England, South Africa, and Brazil and Japan, respectively. All three carried convergent Spike (S) N501Y substitutions, while Beta and Gamma shared substitutions S:K417N/T and S:E484K. It is now appreciated that SARS-CoV-2 variants can be more transmissible [2], differ in pathogenicity [3,4], and/or escape immunity afforded by vaccination or by infection with earlier variants [5,6]. Although the Omicron lineage has displaced Delta [7], new Omicron sub-lineages continue to emerge [8], certain of which harbor concerning mutations such as S:F486V that escape broadly neutralizing antibodies [9,10]. Overall, the pandemic remains largely uncontrolled in virtually all geographies, highlighting the need for ongoing surveillance to inform public health responses.

New SARS-CoV-2 variants are detected and tracked using viral genomic surveillance, an approach wherein the genomes from circulating viruses are sequenced, usually from patient nasal swabs, and data is rapidly shared using global platforms, such as GISAID (www.gisaid.org) [11] or NCBI. These practices collectively enable scientists around the world to track viral evolution across space and time while keeping tabs on mutations that can compromise the utility of diagnostic assays or negatively impact the clinical efficacy of vaccines or monoclonal antibody therapeutics. Effective surveillance relies on prompt viral whole genome sequencing of recent samples across representative geographies and populations and low latency sharing of resulting data. Accordingly, investments in viral whole genome sequencing have increased greatly during the pandemic, with vastly more whole genome sequences of SARS-CoV-2 being shared than any virus in history.

At the outset of the pandemic, SARS-CoV-2 genome sequencing efforts were patchy at best and in many geographies non-existent. Although the U.K. and South Africa set up exemplary viral genomic surveillance at the outset, much of the initial U.S. viral genomic surveillance data came from individuals who generated it at their own initiative, typically relying on discretionary funds, and in some cases, support from private foundations and philanthropies. Once the first variants of concern emerged, the U.S. Centers for Disease Control and the National Institutes of Health began to invest in the development of a SARS-CoV-2 genomic surveillance system. Despite this, many states in the Midwest and Southern U.S. remain poorly represented in the genomic surveillance data [12] even though their residents have been disproportionately impacted by COVID-19 [13–15]. This rural-urban disparity was compounded during the pandemic for racial and ethnic minorities [16,17] who already bore a greater burden of disease [18]. The failure to adequately track emerging variants in a representative fashion may have exacerbated existing disparities in allocation of public health resources, such as rapid antigen tests and monoclonal antibody therapies.

To address this lack of representation, Grambling State University (GSU), Louisiana Tech University (Tech), and Louisiana State University Health Shreveport (LSUHS) established a viral genomic surveillance hub to reach several vulnerable and underserved populations in north Louisiana [19]. Soon after, we partnered with Mercer University School of Medicine (MUSM) in Georgia, Jackson State University (JSU) in Mississippi, as well as with community health centers in each of our areas to increase SARS-CoV-2 genome sequencing volume and representativeness. These efforts resulted in a new viral genomic surveillance network serving a total of 15 counties or parishes across these 3 southern U.S. states (Figure 1).

**Fig 1.**
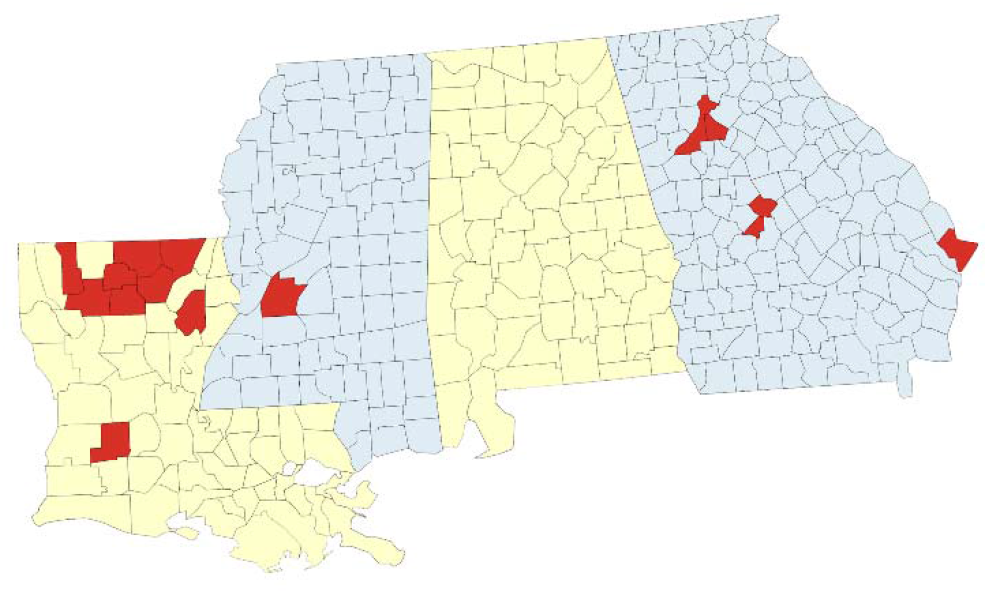
Geographic coverage of SARS-CoV-2 genomic surveillance in Louisiana, Georgia, and Mississippi. Map of the surveillance region with parishes (Louisiana) or counties (Georgia, Mississippi) where at least one specimen was sequenced by the network indicated in red.

Our primary goals were to improve sampling from rural communities and minorities, to empower underrepresented minorities conducting the scientific work of viral genomic surveillance, and to provide individuals who donate samples for viral genome sequencing the opportunity to do so with consent and an understanding of how their samples would be used to track viral variants. The pandemic has underscored the importance of contextual metadata accompanying genomic data. When metadata is incomplete, it hinders comparative analyses and data-driven policymaking [20,21]. Thus, an additional objective was to collect more complete metadata to maximize the usefulness of the shared sequences. Here we used patient metadata to assess the relationships between viral gene copies (as an index of viral load) and variant type, vaccination status, and symptomatology. We report on the results of our work in Louisiana, Georgia, and Mississippi.

## Results and discussion

### Improving representativeness in genomic surveillance

In Louisiana, our six partner clinics collected 405 rapid antigen-positive clinical specimens from the residents of nine parishes (Allen, Bienville, Franklin, Jackson, Lincoln, Morehouse, Ouachita, Union, and Webster) between 22 July 2021 and 12 April 2022 (Fig 1). We worked closely with clinicians at the collection sites to gain informed consent from patients and collected metadata on patient demographics and vaccination status for more than 90% of the specimens (Table 1).

**Table 1.**
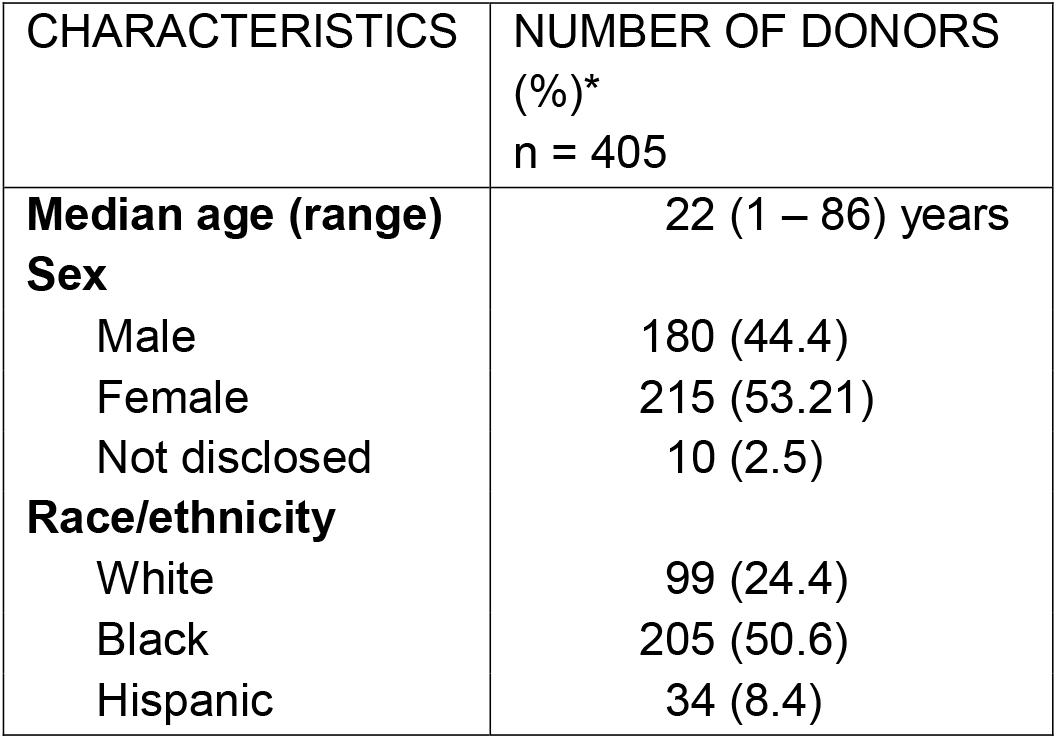
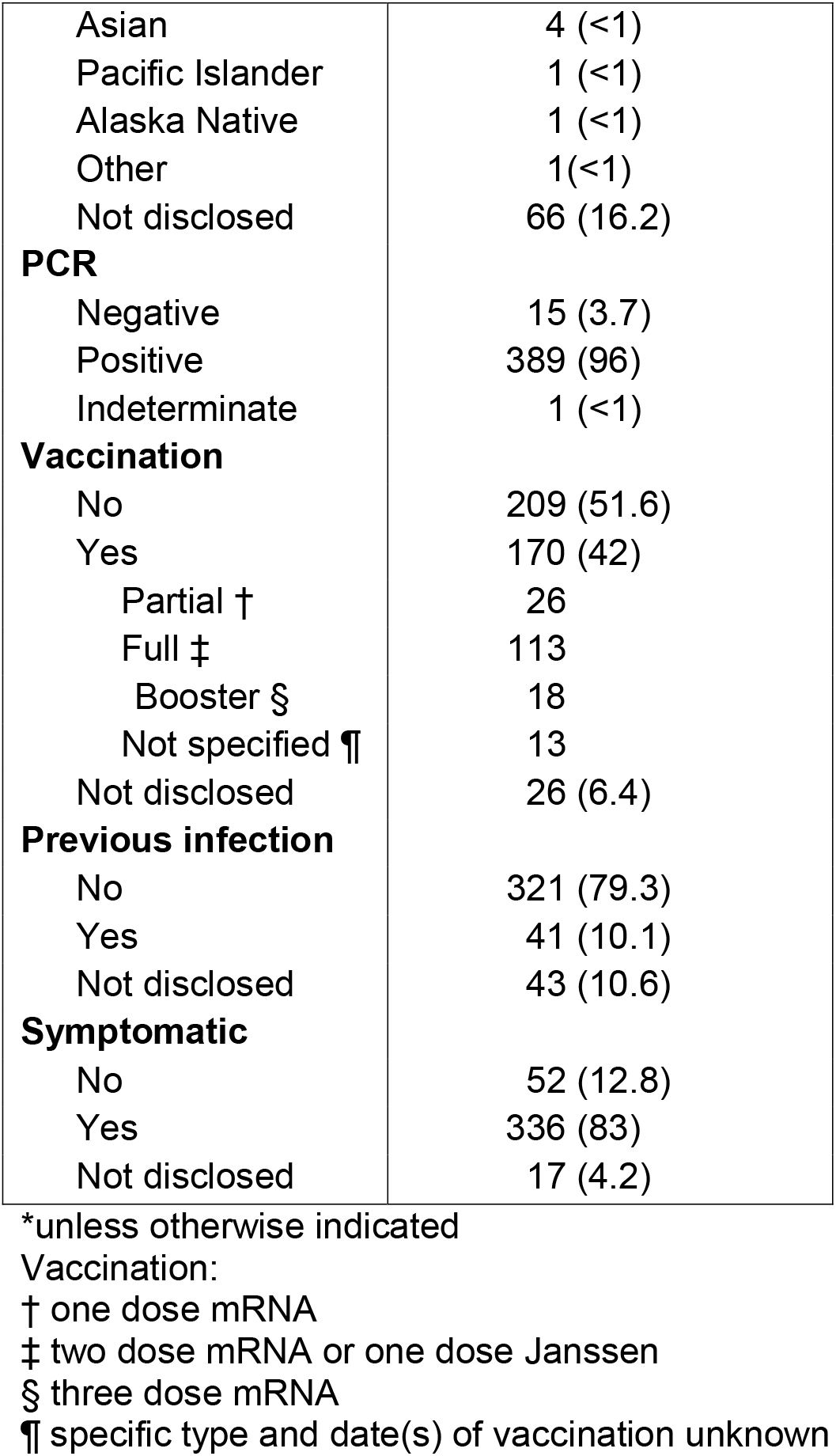
Characteristics of study donors in Louisiana (22 July 2021 – 12 April 2022).

| CHARACTERISTICS | NUMBER OF DONORS<br>(%)*<br>n = 405 |
| --- | --- |
| **Median age (range)** | 22 (1 – 86) years |
| **Sex** |  |
| Male | 180 (44.4) |
| Female | 215 (53.21) |
| Not disclosed | 10 (2.5) |
| **Race/ethnicity** |  |
| White | 99 (24.4) |
| Black | 205 (50.6) |
| Hispanic | 34 (8.4) |
| Asian | 4 (<1) |
| Pacific Islander | 1 (<1) |
| Alaska Native | 1 (<1) |
| Other | 1 (<1) |
| Not disclosed | 66 (16.2) |
| **PCR** |  |
| Negative | 15 (3.7) |
| Positive | 389 (96) |
| Indeterminate | 1 (<1) |
| **Vaccination** |  |
| No | 209 (51.6) |
| Yes | 170 (42) |
| Partial † | 26 |
| Full ‡ | 113 |
| Booster § | 18 |
| Not specified ¶ | 13 |
| Not disclosed | 26 (6.4) |
| **Previous infection** |  |
| No | 321 (79.3) |
| Yes | 41 (10.1) |
| Not disclosed | 43 (10.6) |
| **Symptomatic** |  |
| No | 52 (12.8) |
| Yes | 336 (83) |
| Not disclosed | 17 (4.2) |

\*unless otherwise indicated
Vaccination:
† one dose mRNA
‡ two dose mRNA or one dose Janssen
§ three dose mRNA
¶ specific type and date(s) of vaccination unknown

In these nine parishes, between 24.2% (Ouachita) to 82.9% (Union) of the residents live in rural areas as defined by the U.S. Census Bureau [22]. Rural areas may be especially vulnerable to the pandemic because they tend to have less capacity for diagnostic testing and healthcare services [23,24] while being home to a poorer, older, health-compromised population [25]. Bienville and Franklin parishes in particular ranked in the top 20 of the 300 poorest counties with the highest COVID-19 mortality rates in a recent analysis of 3,200 U.S. counties and the majority of the other parishes in this region were likewise characterized as low income-high mortality [26].

The disproportionate impact of COVID-19 on racial and ethnic minorities has also been well documented [27–29]. More than 60% of our donors self-identified as non-white or as two or more races compared to approximately 41% overall in the sampled parishes. In our view, this oversampling supports equitable genomic surveillance by compensating for known biases in sampling [30] from both rural and minority populations. Notably, 8.4% of our specimens were from Hispanic donors who make up just 3.1% of the northeast Louisiana population but were far more likely to contract COVID-19, accounting for 13.8% of overall cases in the region (up to 60% in some parishes) according to the Louisiana Department of Health. Our partnership with The Health Hut mobile clinic staffed with Spanish-speaking translators improved representativeness in this population. Of the 405 specimens that were collected in Louisiana, 389 had detectable SARS-CoV-2 RNA in the CDC real-time RT-PCR diagnostic assay and underwent sequencing, with genomes ultimately recovered from 272 or 70% of PCR-positive specimens. This represents 1.32% of the 20,617 genomes in GISAID from Louisiana during the same period.

### Accelerating regional genomic surveillance

After successfully establishing relationships with local clinics and developing a pipeline for exchanging specimens and data across our network in Louisiana, we felt confident sharing our model with other peer institutions across the south. Mercer University School of Medicine (MUSM) in Georgia and Jackson State University (JSU) in Mississippi both joined our efforts, building on relationships to engage their own communities in genomic surveillance.

In Georgia, MUSM specimens were collected from Mercer Medicine (MM) Student Health services in Bibb, Chatham, DeKalb, and Fulton counties and an MM community clinic located in Peach county (Fig 1). Roughly 81% of the Georgia specimens were obtained from counties outside of the Atlanta metropolitan area. Approximately 48% of MM donors did not self-identify as white. After testing positive via PCR analysis, specimens were sent to the Center of Excellence for Emerging Viral Threats (CEVT) at LSUHS for Illumina sequencing.

Among other achievements, Mercer’s work identified two emerging subvariants carrying immune escape mutations [31,32]: the first case of BA.2.75 in Georgia, which was only the 40th documented instance of the variant in the U.S., and the sixth case of BQ.1.1 in Georgia, one of the most antibody-evasive variants to date [31]. MM generated 703 SARS-CoV-2 genomes representing 1.8% of the total data in GISAID from Georgia between 13 September 2021, when the first sample was collected by MM to 10 October 2022.

In Mississippi, the JSU Health Services Center served as the collection site (Fig 1). The Center serves as the primary COVID-19 testing site for students, staff, and visitors of the university and supports the surrounding community’s testing and vaccination efforts. The engagement with JSU extended the network’s reach into a predominantly black community. One hundred percent of the 33 samples submitted for sequencing at the CEVT were from non-white donors.

### Building NGS capacity for local genomic surveillance

The specimens collected in Louisiana were initially submitted to the CEVT for sequencing until we built local sequencing capacity at GSU and Tech. By sequencing locally, we achieved a median latency, or turnaround time from sample collection to release on GISAID, of 5 days at GSU and 12 days at Tech when processing contemporary rather than previously stored specimens (Fig 2B). Through this approach, we were able to report the first confirmed case of Omicron in northeast Louisiana, a region comprising 12 parishes and 323,000 residents, within 2 days of sample collection [33]. We then closed the surveillance loop by sharing the variant data along with weekly wastewater surveillance results and other COVID-19-related health information through a dashboard developed for the north Louisiana community (www.nla-health.com/dashboard).

**Fig 2.**
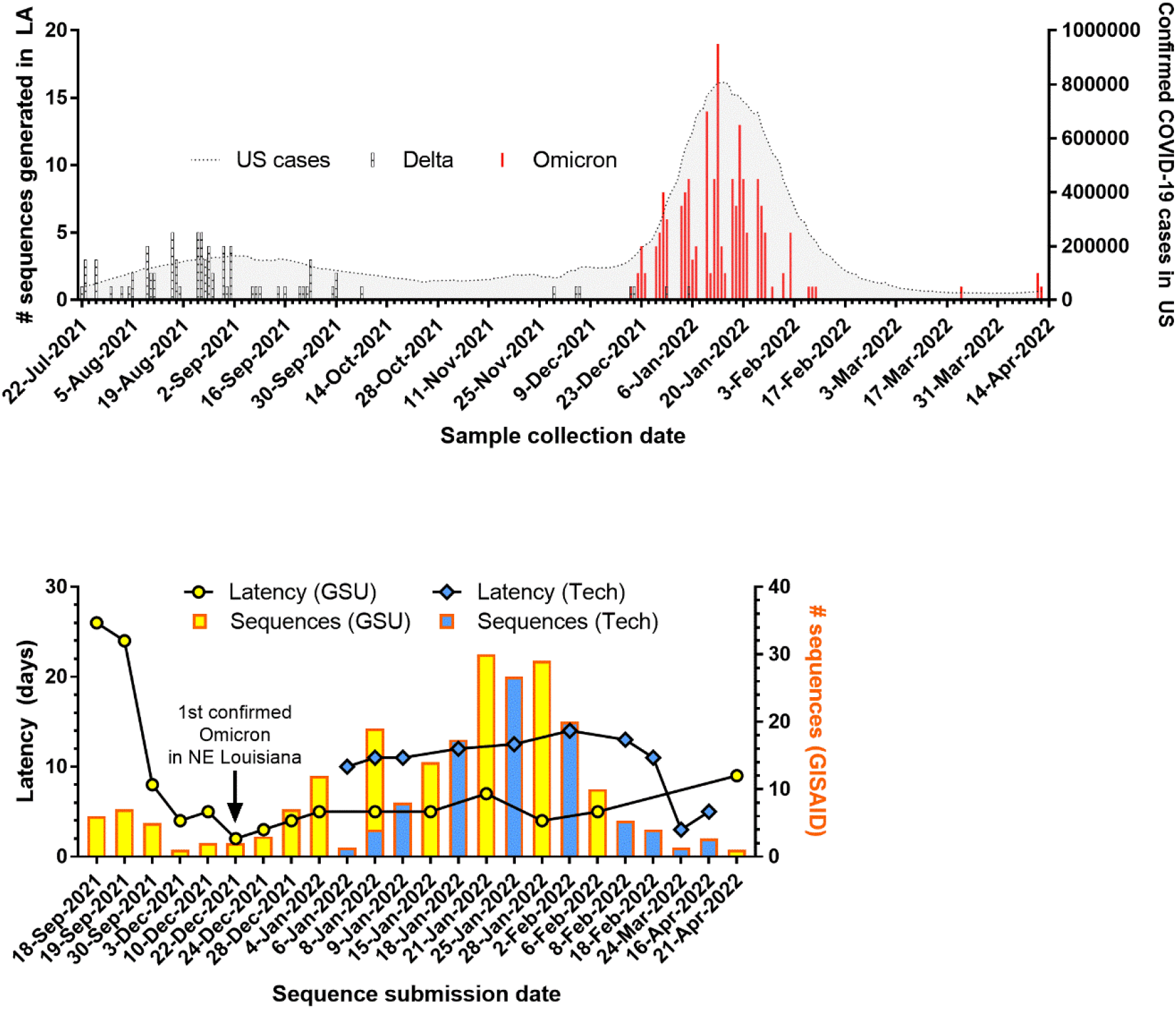
SARS-CoV-2 genomic surveillance in Louisiana. (A) Number of variants sequenced over time in Louisiana overlaid with number of confirmed COVID-19 cases in the U.S. (B) Number of sequences uploaded and latency for new sequencing entities established at Grambling State University (GSU) and Louisiana Tech University (Tech).

The results of sequencing, including comparison of Nanopore to Illumina, investigation of viral genome coverage and copy number, and phylogenetic analyses are described below.

### Nanopore concordance with Illumina

Among the various approaches to whole genome sequencing of viruses [34], multiplex PCR amplification of the viral genome followed by amplicon sequencing is the most widely adopted for SARS-CoV-2 genome sequencing. The second-generation Illumina platform dominates the next generation sequencing (NGS) field, but third-generation Oxford Nanopore technology has attracted attention during the pandemic in large part for its low costs of entry, real-time analysis, and amenability to low-throughput use [35].

We first validated the performance of Nanopore MinION sequencing at GSU and Tech by sequencing a matched subset of 16 SARS-CoV-2 positive specimens using a well-established Illumina workflow at the CEVT. The results confirmed the accuracy of the Nanopore sequencing protocol and analysis pipeline in our setting with 100% consensus sequence identity in 13/16 of the samples and 99.9% sequence identity in 3/16 samples. The one discordant basecall was identical in the three samples: T by Illumina sequencing and G by Nanopore sequencing at 17,259, within the gene encoding the replicase accessory protein NSP13. The reference SARS-CoV-2 genome also contains a G at this position; the T mutation identified by Illumina sequencing results in a glutamic acid rather than an aspartic acid. The E1264D mutation identified in the samples sequenced with the Illumina protocol is well supported. Nanopore basecalling accuracy decreases in homopolymeric tracts [36] but the reason for the discordance observed in this region is unclear.

Importantly, all sequences were 100% concordant for the Pango lineages and GISAID clade assignments. The 16 samples sequenced by both protocols spanned a wide range of C_q_ values (15.6 – 32.4) but almost all samples, Nanopore or Illumina, generated > 98% genome coverage. Illumina sequencing resulted in an average of 285 bases more sequence data than Nanopore, but this was mainly confined to the extreme 5’ and 3’ ends of the genome.

### Viral genome coverage

Plotting the breadth of coverage for Nanopore sequencing reveals a threshold of approximately C_q_ 30 for reliably recovering a full genome in our setting and an upper limit of approximately C_q_ 35 (Fig 3). These coverage thresholds are congruent with values from the literature of different NGS methodologies for SARS-CoV-2 genome sequencing [37], although some groups using the Nanopore platform have reported high coverage consensus genome sequences from specimens with C_q_ values up to 39 [38]. Variability in primer design and amplification parameters, library preparation method, and sequencing run time can influence results. For example, longer amplicons reduce the likelihood of primer interactions [39] and primer mismatches due to mutations [40], generally outperforming shorter amplicons [41,42]. Shorter amplicons, however, can recover more complete genomes from specimens where the viral load is low or the RNA is degraded [43,44].

**Fig 3.**
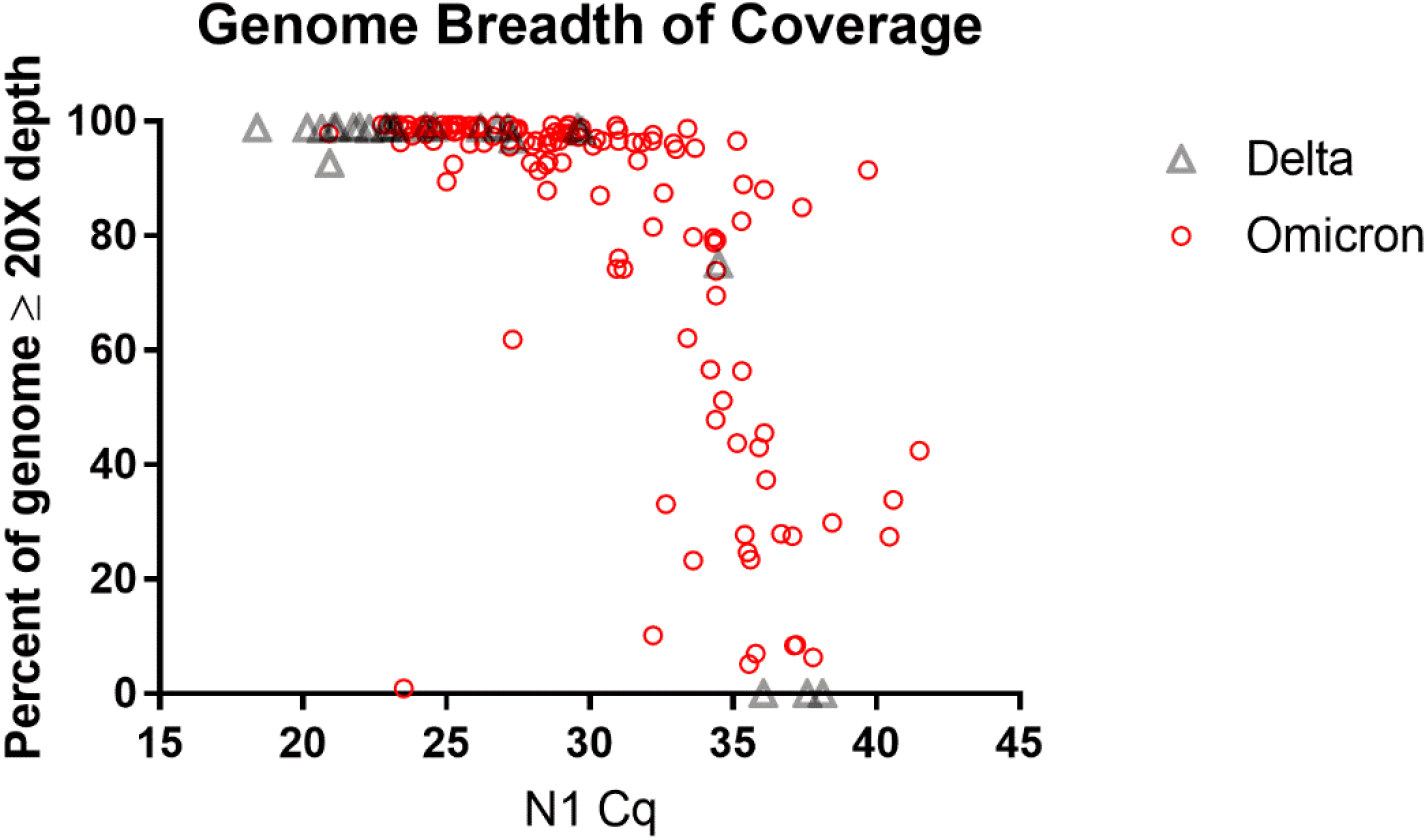
Decreasing genome breadth of coverage in samples with high C_q_ values. Higher C_q_ values were observed in Omicron (n = 134) than in Delta (n = 27). All but one Delta specimen was amplified for 32 cycles and all Omicron specimens were amplified for 35 – 40 cycles.

Specimens carrying low viral loads can have uneven genome coverage due to poor amplification of the less efficient amplicons, which can be addressed to some extent by optimizing the PCR and sequencing protocols. Lagerborg *et al*. found that increasing PCR cycles from 35 to 40 resulted in more uniform coverage for Illumina-based ARTIC sequencing of low viral load specimens while still avoiding PCR artifacts [44]. Compared to the Nanopore rapid barcoding kit used for library preparation in our study, the more labor-intensive Nanopore ligation sequencing kit can yield greater sequencing depth and better (*de novo*) assembly of the SARS-CoV-2 genome [45]. Of course, longer run times can produce more mapped reads to help recover complete genomes.

### Viral gene copies

The fast-spreading Omicron variant displaced the Delta variant in our sampling with little overlap in prevalence (Fig 2A). If we infer that low coverage, unclassified sequences are the predominant variant at the time of sample collection, we find that the mean C_q_ value of Omicron specimens (29.87, 95% CI 29.07 – 30.67) was significantly higher than that of Delta specimens (25.55, 95% CI 23.38 – 27.72). This corroborates early reports of lower viral gene copy number determined by RT-qPCR [46,47] or lower infectious viral load determined by focus forming assay [48] in Omicron infections compared to Delta infections.

One possible explanation for the observed difference in gene copy number/imputed viral load is that during the initial Omicron surge, a greater proportion of the population may have acquired some degree of immunity through vaccination or prior infection. Some reports support the idea of attenuated viral load with vaccination [49–51] while others suggest no significant difference between unvaccinated and vaccinated individuals [52–54].

Having collected vaccination and prior infection metadata from 379 and 362 of the 405 donors respectively, we compared immune history with viral gene copy numbers. We found that the mean C_q_ value of unvaccinated individuals with no previous infection (n = 82) was 1.06 cycles lower than that of fully vaccinated individuals with or without a previous infection (n = 81), but this difference was not statistically significant (p = 0.218) (Fig 4A).

**Fig 4.**
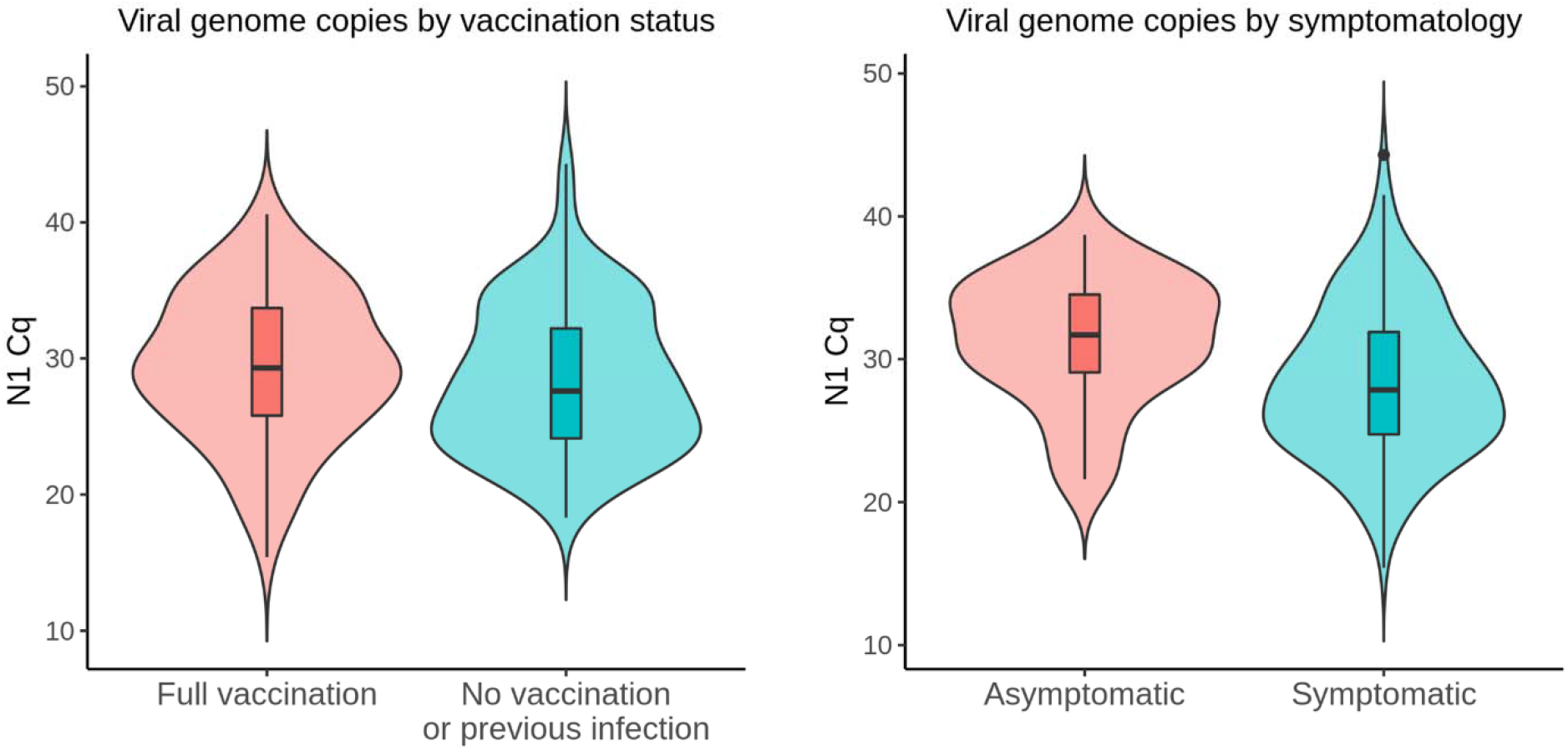
Viral gene copies by vaccination status and symptomatology. (A) No statistically significant differences (p = 0.218) in viral gene copies were observed between unvaccinated donors without a previous infection (n = 82) and vaccinated donors with or without a previous infection (n = 81). (B) Viral gene copies were lower (p = 0.0039) in asymptomatic donors (n = 30) compared to symptomatic donors (n = 164).

Several groups have found no correlation between RT-qPCR quantification of viral genes and symptomatology [55–58]. Two studies reported higher C_t_ values in asymptomatic individuals (n = 5 or 3) compared to symptomatic individuals (n = 14 or 9) but the sample sizes were limited [59,60]. In a larger study, asymptomatic individuals (n = 2179) exhibited higher C_t_ values compared to symptomatic individuals (n = 739), but the authors concluded that the statistically significant difference of 0.71 cycles was not clinically meaningful [61]. In our dataset, the mean C_q_ value of asymptomatic donors (31.36, 95% CI 29.7 – 33.01; n = 30) was nearly 3 cycles higher than that of symptomatic donors (28.38, 95% CI 27.56 – 29.19; n = 164) (Fig 4B).

The asymptomatic donors in this study were mostly college students who tested positive during baseline surveillance on the GSU campus, so the age distribution is highly skewed compared to symptomatic donors. There are conflicting reports on the relationship between patient age and viral load, but one of the largest cross-sectional studies found that viral load tends to increase with age [62]. To account for a possible age effect, we compared asymptomatic donors who were age 30 or younger (n = 28) with symptomatic donors in the same age group (n = 93) and found that the asymptomatic group still had lower mean viral gene copy number (31.78, 95% CI 30.17 – 33.39 vs 28.55, 95% CI 27.39 – 29.7).

Only C_q_ values derived at GSU were used in these analyses to avoid potential confounding by interlaboratory variation in PCR assays [63]. These analyses should be considered carefully within the limitations of our study which include a relatively small sample size and lack of longitudinal information on peak copy number, and also within the overarching context that copy number is an imprecise index of viral load [64].

### Phylogenetic analysis of northeast Louisiana SARS-CoV-2 genomes

We performed independent phylogenetic analysis for Delta and BA.1.x SARS-CoV-2 viral populations from Louisiana and Georgia in the context of representative global samples. Time-scaled maximum likelihood phylogenies revealed that Delta and BA.1.x isolates from Louisiana and Georgia are interspersed throughout the global population suggesting multiple independent introductions (Fig 5). Often, these introductions resolve into distinct subclades indicative of ongoing transmission within a new location. Further, putative introductions were spatially diverse and included Europe and South America as well as other states throughout the US. These local epidemics were nested within the larger North American epidemic for both Delta and BA.1.x waves.

**Fig 5.**
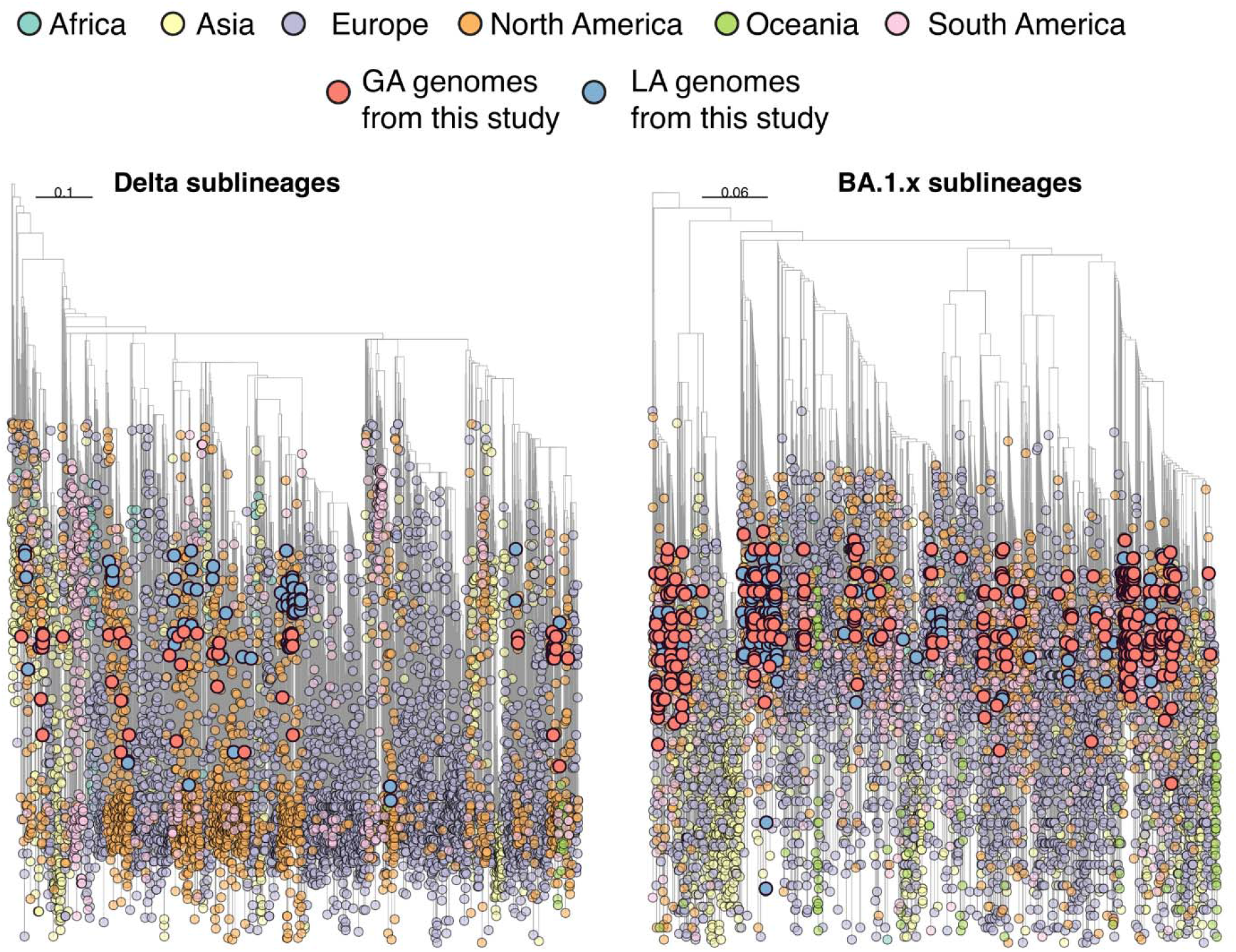
Time-resolved Maximum Likelihood (ML) phylogenies of Delta and Omicron (BA.1.x) sublineages. The genomes generated in this study are colored in red for Georgia and blue for Louisiana. Tree tips are colored according to their location (continent) and the color legend is on the top of the figure.

## Conclusion

### NGS at scale

Nanopore sequencing has been used for real-time genomic surveillance during outbreaks of Ebola in West Africa [65], Zika [66] and Yellow Fever [67] in Brazil, and Lassa fever in Nigeria [68]. During the COVID-19 pandemic, laboratories around the world applied Nanopore sequencing to rapidly share SARS-CoV-2 genomic data. The Nanopore platform is especially suitable for smaller institutions due to its low capital and operational cost, accessible infrastructure requirements, and high degree of scalability. A modestly equipped molecular biology laboratory with the appropriate biosafety containment can implement the Nanopore rapid barcoding workflow to go from swab to data for less than $3500 USD at the time of this writing. The operational cost (excluding RNA extraction) has been estimated at £18.91 per ligated barcode for 12 barcodes on a MinION flow cell when using a wash kit to remove the previous library [69]. This corresponds with our cost analysis (*including* RNA extraction) of $20 – $25 USD per tagmented barcode for 12 barcodes when the flow cell is washed and reused.

Working with NGS data can demand significant computational resources and bioinformatics expertise, but Nanopore-based SARS-CoV-2 sequencing can be performed on a consumer-class PC using prepared workflows including point-and-click pipelines in the cloud. Although higher throughput laboratories can multiplex 96 barcodes for less than £10 per barcode [69], the platform does allow small-volume laboratories to cost-effectively conduct genomic surveillance in areas of low sequencing capacity.

The democratization of genomics is nowhere better demonstrated than in Africa where technological advances in NGS coupled with major investments by Africa CDC and partners rapidly expanded genomic surveillance across the continent [70]. The efforts in Africa highlight some parallels from our experience in the rural Southern U.S., that local sequencing programs can close surveillance blind spots while significantly decreasing turnaround time. This is exemplified by our sequencing the first confirmed case of Omicron in northeast Louisiana and detecting the emergence of BA.2.75 and BQ.1.1 in Georgia.

### Collaborative community-centered genomic surveillance networks

As part of a broader initiative supported by the Rockefeller Foundation’s Pandemic Prevention Institute, we established a genomic surveillance network in our community focused on equitable sequencing among a more vulnerable, marginalized population. We formed collaborations between R1 institutions, primarily undergraduate universities, a graduate medical school, Historically Black Colleges and Universities, and community health centers to collect and sequence respiratory specimens from patients in rural Louisiana, Georgia, and Mississippi. These three states rank 41st, 43rd, and 45th respectively among U.S. states in the percentage of cumulative cases sequenced [12]. With a modest investment from the Rockefeller Foundation and internal support from our institutions, we increased sequencing volume from these states so that sequenced samples better represent the diversity of the U.S. population. We collected enriched metadata including vaccination status and county/parish-level geographical location, which is particularly important when emerging variants such as BA.2.75 and BQ.1.1 are detected. In both the GSU and Tech initiatives, health care workers at partnering clinics briefly educated patients about how viral genomic surveillance is used to track emerging variants and provided patients with the opportunity to provide consent for their positive sample to be used for whole viral genome sequencing.

Rural residents face significant barriers rooted in geographic isolation, scarcity of services, lower socioeconomic status and other social determinants of health, and even cultural constraints that impede them from getting the healthcare that they need [71]. During the pandemic, this disparity manifested as disproportionately high mortality rates in rural America, particularly among minorities [72]. In rural areas, so-called regional universities represent essential infrastructure that can provide not only educational opportunities but also access to testing and vaccination. These universities are uniquely positioned to gain trust and encourage participation in genomic surveillance because they often function as community anchors, vital to the local economy, civics, and culture [73]. A viral genomic surveillance strategy that leverages these strong community ties also offers a much-needed on-ramp for students from diverse groups to gain genomics research experience that will prepare them to combat misinformation in their own communities and participate in the genomics workforce.

Our work demonstrates the impact that universities have within their local communities, the power of the relationships that can be built between academia and clinics, and the positive influence of these partnerships on public health. We propose that these types of nimble, local networks should continue to play an important role in national efforts to modernize viral genomic surveillance of common respiratory illnesses by filling in critical gaps in genomic surveillance coverage, consent, *and* participation.

## Methods

### Specimen collection

Patients testing positive (*n* = 405) on the BinaxNOW rapid antigen test (Abbott Laboratories; Abbott Park, IL) at the Foster Johnson Health Center in Grambling State University (Grambling, LA), The Health Hut (Ruston, LA), Louisiana Tech University (Ruston, LA), Michael Brooks Family Clinic (Ruston, LA), Minden Family Care Center (Minden, LA), and Serenity Springs Specialty Hospital (Ruston, LA) between 23 August 2021 and 12 April 2022 were recruited through an informed consent process. Remnant anterior nasal swabs used in the rapid test (The Health Hut, Michael Brooks Family Clinic, Louisiana Tech University, Minden Family Care Center, Serenity Springs Specialty Hospital) or freshly collected mid-turbinate nasal swabs (Foster Johnson Health Center) were preserved in 3 mL of viral transport media (VTM) at 4 °C and later in the study in 1 mL of DNA/RNA Shield at room temperature. The work at GSU was conducted under 45 CFR 46.102(l)(2) as a public health surveillance activity and exempt from IRB review. The work at Tech was conducted under IRB protocol HUC 21-106. LDT COVID-19 IMD_A_ specimen collection at Mercer University (Macon, GA) was conducted under IRB protocol H2110206 using previously collected anterior nasopharyngeal swabs or mid-turbinate nasal swabs, and the samples were sequenced at LSUHS CEVT. Specimen collection at Jackson State University (Jackson, MS) was conducted under IRB protocol 0097-22 using either anterior nasal swabs or mid-turbinate nasal swabs, and the samples were sequenced at LSUHS CEVT. Sequencing at LSUHS CEVT was conducted under IRB protocol # STUDY00001445.

### Illumina sequencing (LSUHS CEVT)

Total RNA was isolated from VTM within 72 hours of specimen collection using the Omega Bio-Tek Viral RNA Xpress kit (Norcross, GA) on an automated extraction platform. Presence of SARS-CoV-2 RNA was determined by using CDC primers and probes with LunaScript RT Supermix Kit (NEB) run on BioRad (Hercules, CA) C1000 real-time PCR machines. cDNA, PCR amplification, and library preparation was performed using the NEBNext ARTIC SARS-CoV-2 FS kit (NEB) according to the manufacturer’s instructions. Libraries were quantified using Invitrogen Qubit BR dsDNA kit and size determined on an Agilent Tapestation 2200 using high sensitivity d1000 tapes (Santa Clara, CA). Libraries were loaded at 10.5 pM on an Illumina (San Diego, CA) MiSeq version 2, 300 cycle kit on an Illumina MiSeq instrument. Consensus sequences were generated with an in-house data analysis pipeline using bwa, samtools, and ivar.

### Nanopore sequencing (GSU and Tech)

Total RNA was extracted from VTM or DNA/RNA Shield within 48 hours of specimen collection via the QIAmp Viral RNA Minikit (Qiagen; Germantown, MD) or Zymo Quick-DNA/RNA Viral MagBead Kit (Zymo Research; Irvine, CA) according to the manufacturers’ protocols. Briefly, 500 uL of VTM was centrifuged at 5,000 g for 10 minutes to pellet cells and debris and either 140 uL of supernatant was extracted in 560 uL of Buffer AVL containing 560 ug of carrier RNA (QIAmp) or 200 uL of supernatant was extracted in 400 uL of Viral DNA/RNA Buffer followed by addition of 10 uL MagBinding Beads (Zymo). After washing the spin column or beads with the manufacturers’ respective wash buffers, total RNA was eluted from the column in 60 uL of Buffer AVE (QIAmp) or from the beads in 30 uL of DNase/RNase-free water (Zymo) and immediately used for RT-qPCR and library preparation. The presence of SARS-CoV-2 was confirmed using the SARS-CoV-2 Research Use Only qPCR Primer and Probe Kit (Integrated DNA Technologies, IDT; Coralville, IA) and qScript XLT One-Step RT-qPCR ToughMix (Quantabio; Beverly, MA) according to the CDC 2019-nCoV Real-Time RT-PCR Diagnostic Panel procedure.

The sequencing library was prepared using the 1200 bp amplicon “midnight” primer set (versions 1 or 2) [72] following the protocol by Freed and Silander [38]. Briefly, the RNA was reverse transcribed using the LunaScript RT Supermix Kit (New England BioLabs, NEB; Ipswich, MA). Tiled 1200 bp amplicons were generated using midnight primers (IDT) with Q5 Hot Start High-Fidelity 2X Master Mix (NEB) for 32 cycles (LA-GSU1 to LA-GSU19) or up to 40 cycles (LA-GSU20 to LA-GSU148 and LA-TECH1 to LA-TECH68) of multiplex PCR amplification. The two overlapping amplicon pools for each specimen were combined and quantified using the Qubit dsDNA HS Assay Kit (Invitrogen; Carlsbad, CA). A total of 75 – 150 ng of PCR product per specimen was barcoded using the Nanopore Rapid Barcoding Kits SQK-RBK004 or SQK-RBK110.96 (Oxford Nanopore Technologies, ONT; Oxford, UK). The barcoded samples were pooled and purified using AMPure XP beads at a 1:1 ratio of sample:beads (Beckman Coulter; Brea, CA). The final libraries (350 – 800 ng DNA) were sequenced on an R9.4.1 flow cell using a MinION Mk1B or Mk1C device, basecalled in real-time via MinKNOW, and analyzed using the wf-artic workflow (medaka, minimap2, bcftools, samtools, nextclade, artic, pangolin) according to the ONT protocol.

### Phylogenetic analysis

SARS-CoV-2 genomes were downloaded from GISAID (www.gisaid.org) excluding the low coverage and incomplete records up to April 30th 2022, and subsampling carried out using subsampler to obtain a representative dataset [73]. Subsampling 4,199,772 (Delta sublineages) and 2,895,902 (BA.1.x sublineages) genomic sequences resulted in 5,882 (Delta sublineages) and 6,843 (BA.1.x sublineages) total sequences. Viral sequences were aligned using ViralMSA with default parameters [74] using Wuhan-1 (MN908947.3) reference and then manually curated with Aliview to designate the start site and trim terminal regions [75]. A maximum likelihood (ML) phylogeny was inferred using IQTREE v1.6.10 with the best-fit nucleotide substitution model identified by the Model Finder function [76]. Statistical support for nodes was determined using 1,000 bootstrap replicates. TreeTime [77] was used to transform this ML tree topologies into a dated tree using a constant mean rate of 8.0 × 10^-4^ nucleotide substitutions per site per year, after the exclusion of outlier sequences. The phylogeny was visualized using the *ggtree* package in RStudio with tips colored by collection source (continent).

### Statistical analysis

Statistical analysis was performed using Prism version 7.05 (GraphPad; San Diego, CA). Continuous variables between two groups were tested for normal distribution (D’Agostino & Pearson test) and homogeneity of variance (F test) then compared using an independent t-test with two-tails and alpha set to 0.05.

## Data availability

The 1,053 complete SARS-CoV-2 genomes that have been sequenced to date by this initiative using Illumina or Nanopore sequencing are available on GISAID and accessible at doi.org/10.55876/gis8.221011tg. The phylogenetic analyses are based on sequences available on GISAID and accessible at doi.org/10.55876/gis8.220901cu.

## Acknowledgements

The authors deeply thank the healthcare professionals at Foster Johnson Health Center (Dawn Holmes), The Health Hut (Jackie White, Chelsea Streets), Michael Brooks Family Clinic (Natalie Wise, Taylor Schutzman, Whitney Walker), Minden Family Care Center (Beau Burton), Serenity Springs Specialty Hospital (Tonya Monk), Mercer Medicine (Certified Medical Assistants), and Jackson State University’s Health Services Center (Kimberly Bullock) for recruiting patients, gaining informed consent, and/or collecting the clinical specimens. We thank Maarten Van Diest for assistance with data curation. We are grateful to the Louisiana Department of Health for their efforts to facilitate enhanced viral genomic surveillance as a public health activity. We gratefully acknowledge all data contributors, i.e., the Authors and their Originating laboratories responsible for obtaining the specimens, and their Submitting laboratories for generating the genetic sequence and metadata and sharing via the GISAID Initiative, on which this research is based. The full acknowledgement table can be found in Supplemental Table 1 and 2. We gratefully acknowledge the contribution of the GISAID Data Curation Team to enable rapid sharing of genomic data. We acknowledge the Louisiana Biomedical Research Network and its Bioinformatics, Biostatistics, and Computational Biology Core for providing computational resources.

## Financial disclosure for submission system

This work was supported by a grant from The Rockefeller Foundation (2021 HTH 010) to PYK, JJN, JPK. PYK acknowledges the Office of the Provost at GSU for funding. JJN acknowledges the School of Biological Sciences at Tech for funding. JPK also acknowledges support from National Institutes of Health grants no. P20GM121307-04S1, P20GM121307 and P20GM134974 and a COVID-19 Fast Grant (no. 2239) from Emergent Ventures, an initiative of the Mercatus Center at George Mason University. The funders had no role in study design, data collection and interpretation, or the decision to submit the work for publication.

## Author contributions for submission system

PYK: conceptualization, formal analysis, funding acquisition, investigation, project administration, writing - original draft preparation, writing - review & editing

AYK: investigation, supervision, writing - review & editing

JJN: conceptualization, funding acquisition, investigation, project administration, writing - review & editing

EC: formal analysis, visualization, writing - review & editing

TCB: software, resources, writing - review & editing

GW: data curation, project administration.

JC: data curation, project administration

JFS: data curation

KHD: data curation, project administration, visualization

IVO: data curation, investigation, project administration, writing - review & editing

OA: investigation

SJ: investigation, resources

ONU: data curation, investigation, resources

RJM: project administration, supervision

VLM: data curation, investigation, resources, software, validation

MPH: project administration, writing - review & editing

PJH: project administration, supervision

MF: data curation, investigation

LV: data curation, investigation

JT: investigation

TA: formal analysis, writing - review & editing

KQ: data curation, investigation, project administration, writing - review & editing

JPK: conceptualization, data curation, funding acquisition, investigation, project administration, writing - review & editing

## Notes

### Competing Interest Statement

The authors have declared no competing interest.

### Summary of Updates

Moved figures next to figure captions for readability. Section on 'Accelerating Regional Genomic Surveillance' updated to clarify emerging variants identified in Georgia.

